# Knockout of AMPA receptor binding protein Neuron-Specific Gene 2 (NSG2) enhances associative learning and cognitive flexibility

**DOI:** 10.1101/2024.02.23.581648

**Authors:** Amber J. Zimmerman, Antonio Serrano-Rodriguez, Sandy J. Wilson, David N. Linsenbardt, Jonathan L. Brigman, Jason P. Weick

## Abstract

The vast majority of gene mutations and/or gene knockouts result in either no observable changes, or significant deficits in molecular, cellular, or organismal function. However, in a small number of cases, mutant animal models display enhancements in specific behaviors such as learning and memory. To date, most gene deletions shown to enhance cognitive ability generally affect a limited number of pathways such as NMDA receptor- and translation-dependent plasticity, or GABA receptor- and potassium channel-mediated inhibition. While endolysosomal trafficking of AMPA receptors is a critical mediator of synaptic plasticity, mutations in genes that affect AMPAR trafficking either have no effect or are deleterious for synaptic plasticity, learning and memory. NSG2 is one of the three-member family of Neuron-specific genes (NSG1-3), which have been shown to regulate endolysosomal trafficking of a number of proteins critical for neuronal function, including AMPAR subunits (GluA1-2). Based on these findings and the largely universal expression throughout mammalian brain, we predicted that genetic knockout of NSG2 would result in significant impairments across multiple behavioral modalities including motor, affective, and learning/memory paradigms. However, in the current study we show that loss of NSG2 had highly selective effects on associative learning and memory, leaving motor and affective behaviors intact. For instance, NSG2 KO animals performed equivalent to wild-type C57Bl/6n mice on rotarod and Catwalk motor tasks, and did not display alterations in anxiety-like behavior on open field and elevated zero maze tasks. However, NSG2 KO animals demonstrated enhanced recall in the Morris water maze, accelerated reversal learning in a touch-screen task, and accelerated acquisition and recall on a Trace Fear Conditioning task. Together, these data point to a specific involvement of NSG2 on multiple types of associative learning, and expand the repertoire of pathways that can be targeted for cognitive enhancement.

## INTRODUCTION

The Neuron-specific gene (NSG) family consists of three (NSG1-3), brain-enriched proteins that regulate proteolytic processing and trafficking of multiple cargos through the secretory and endolysosomal system in neurons. Previous studies have shown that Neuron-specific gene 1 (NSG1; NEEP21) and NSG3 (Calcyon, Caly) regulate diverse cargos including GPCRs (Debaigt et al., 2004), transporters (Muthusamy et al., 2012) and ligand-gated ion channels (Davidson et al., 2009; Steiner et al., 2005). In addition, NSG1 and NSG3 regulate proteolytic processing of Neuregulin-1 and Amyloid Precursor Protein (APP), genes that play critical roles in Schizophrenia and Alzheimer’s Disease, respectively (Muthusamy et al., 2012; Norstrom et al., 2010; Plooster et al., 2021; Yin et al., 2015). However, the role of NSGs in synaptic plasticity via changes in AMPAR trafficking is the most well-characterized. Down-regulation of NSG1 impedes endosomal recycling of GluA1 and GluA2 in hippocampal neurons treated with NMDA (Steiner et al., 2005), and disruption of NSG1 function via dominant negative peptide expression causes reductions in Long-Term Potentiation (LTP) in organotypic hippocampal slices (Alberi et al., 2005). In contrast, NSG3/Calcyon is critical for clathrin-mediated endocytosis of AMPARs, where overexpression reduces AMPAR surface expression and knockout impairs long term depression (LTD; Xiao et al. 2006; Kruusmagi et al. 2007; Davidson et al. 2009; Vazdarjanova et al. 2011).

Despite their opposing roles in synaptic function, animal models with alterations in NSG1 and NSG3 expression do not display predictable and complementary behavioral changes. Our previous work found that global knockout of NSG1 caused mild alterations in motor coordination, significant *increases* in anxiety-like behavior in elevated mazes, but no change in hippocampal- and amygdala-dependent learning and memory (Austin et al., 2022). In contrast, behavioral studies on mice overexpressing (OE) NSG3 found *reductions* in anxiety, where animals spent more time in the light areas of a light-dark box and open areas of the elevated plus maze (Trantham-Davidson et al., 2008). Furthermore, while NSG1 KO animals displayed normal learning and memory, NSG3 OE caused significant perseveration during reversal learning of the Morris Water Maze task, as well as during the extinction phase of context-dependent fear conditioning (Vazdarjanova et al., 2011). Thus, the roles of NSG proteins on behavior appear to be significantly more complex than can be predicted from their demonstrated function at excitatory synapses.

Neuron-specific gene 2 (NSG2) is the least well-characterized member of the NSG family, which arose specifically during the evolution of the vertebrate clade (Muthusamy et al., 2009) suggesting a potential role in more advanced cognitive, affective and/or motivated behaviors. Like other family members, NSG2 plays a role in endolysosomal trafficking and synaptic function. And while the role of NSG2 in specific neural circuits and during synaptic plasticity remains to be determined, previous studies have demonstrated that NSG2 *promotes* excitatory synaptic transmission, where NSG2 OE increased the amplitude of miniature excitatory postsynaptic currents (mEPSCs), while CRISPR-mediated KO reduced mEPSC frequency (Chander et al., 2019). Like NSG1 and NSG3, NSG2 displays a broad distribution throughout excitatory cortical, subcortical, and hippocampal neurons (Barford et al., 2017). Interestingly however, its highest expression is found in multiple regions of the basal ganglia, including the dopamine receptor-expressing cells of the striatum and nucleus accumbens (Genotype-Tissue Expression (GTEx) release version 8; dbGaP Accession phs000424.v8.p2), which is unique among the family members and suggests a possible role in motivated behaviors and habit learning. However, to date nothing is known regarding the impact of NSG2 on any behavioral paradigms. Thus, we surveyed the impact of loss of NSG2 across motor, affective, and cognitive domains. Surprisingly, we found that NSG2 KO animals displayed normal behavior across multiple motor and anxiety-related domains, but displayed *enhanced* associative learning and memory across striatal-, amygdala-, and hippocampal-dependent tasks. To our knowledge, this is the first report of enhanced learning following loss of any NSG family members as well as for proteins that primarily regulate endolysosomal trafficking of AMPARs.

## RESULTS

### Validation of NSG2 KO mice

We first verified the loss of NSG2 protein in null animals. Figure 1A displays a western blot of cortical samples from five wild-type (wt) and five NSG2 knockout (KO) animals that was probed for both NSG1 (upper blot, upper bands) and anti-NSG2 (upper blot, lower bands). WT samples displayed robust staining for both proteins at the appropriate molecular weights (Fig. 1A, left five lanes; ∼21 kDa and 19kDa, respectively), while NSG2 KO samples showed only positive staining for NSG1, indicating a specific loss of NSG2 expression (Fig. 1A, right five lanes). We also ran a separate blot for NSG3 (Fig. 1A, middle blot) to determine whether any compensatory changes may occur between family members. GAPDH signal (lower panel) was used as a loading control to quantify relative band volume across groups (Fig. 1A, lower blot). Pooled data show no significant changes in either NSG1 or NSG3 expression in NSG2 KO animals compared to wt controls (Figure 1B; p<0.05). To demonstrate loss of NSG2 in KO animals occurred throughout the brain, we probed parasagittal sections of wt (Fig. 1C) and KO animals (Fig. 1D). Wild-type animals displayed robust expression of both NSG1 (green) and NSG2 (red) throughout most brain regions, with unique expression of NSG1 in cerebellar Purkinje neurons, consistent with previous findings (Barford et al., 2017). In contrast, sections from NSG2 KO animals showed complete absence of specific staining (Fig. 1D, upper right panel). Thus, we were confident that the CRISPR gene targeting strategy specifically eliminated NSG2 protein expression, while leaving expression of other NSG family members intact throughout the brain.

**Figure 1.**
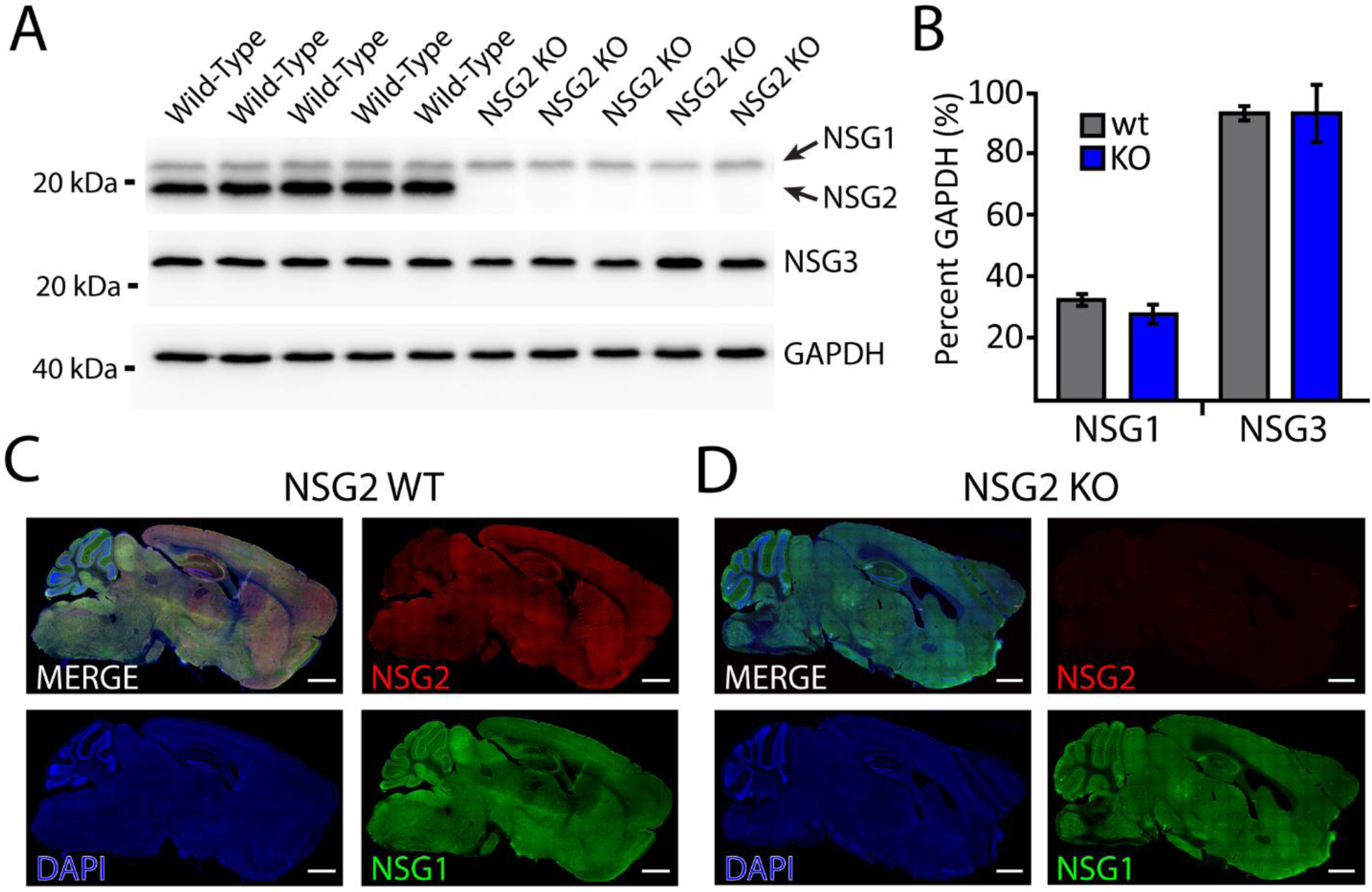
Validation of brain-wide NSG2 knockout in null animals. (A) Western blots of wt C57Bl/6n animals (lanes 1-5) and NSG2 KO animals (lanes 6-10) used for behavioral studies demonstrates complete lack of NSG2 protein in KO animals. (B) Bar graphs of WB band intensity normalized to GAPDH revealed that both wt and KO animals demonstrate normal expression of NSG1 and NSG3. (C-D) Immunohistochemical staining of NSG1 (green), NSG2 (red), and DAPI (blue) on parasagittal brain sections from a wt (C) and KO animal (D). Note specific loss of NSG2 staining in KO section (D). Specificity of NSG1 vs. NSG2 staining is confirmed by robust cerebellar staining of NSG1 antibody in wt brain section (C, lower right panel) while NSG2 signal is relatively low in cerebellum (C, upper right panel).

### NSG2 KO animals show normal motor function

We previously found significant motor coordination deficits in NSG1 KO animals, likely due to its relatively high expression in cerebellum compared with other NSG family members (Austin et al., 2022). We first used the accelerating rotarod test in NSG2 KO animals as a measure of gross motor impairment as well as motor learning. Consistent with previous reports (Austin et al., 2022) we found a significant main effect of trial across groups (F_(4, 88)_=9.27, p<0.0001), indicating that both wt and NSG2 KO animals learned to remain on the beam at higher rotation rates (Figure 2A). However, both groups performed equally well, with no significant difference in the latency to fall between wt and KO mice across all five trials (F(4, 88)=0.8997, p = 0.47). Thus, NSG2 KO animals demonstrated equivalent motor learning with that of wt animals. We next used the Catwalk XT apparatus to determine whether loss of NSG2 would affect overall gait and locomotion. Figure 2B illustrates representative images of step patterns from wt (upper panel) and NSG2 KO (lower panel) animals along with pressure distribution plots of individual footfalls. Figure 2C illustrates a heat map of 174 individual parameters measured across the four paws as well as analysis of coordinated movements, where hotter colors indicate increased levels of particular traits, while cooler colors indicate decreased levels. A volcano plot of these data illustrate that while several parameters met the criteria for significance based on individual t-tests (p<0.05, red line), no traits survived False Discovery Rate analysis to control for the large number of measurements (p=0.0002, dashed cyan line). Thus, loss of NSG2 does not significantly affect motor learning or coordination.

**Figure 2:**
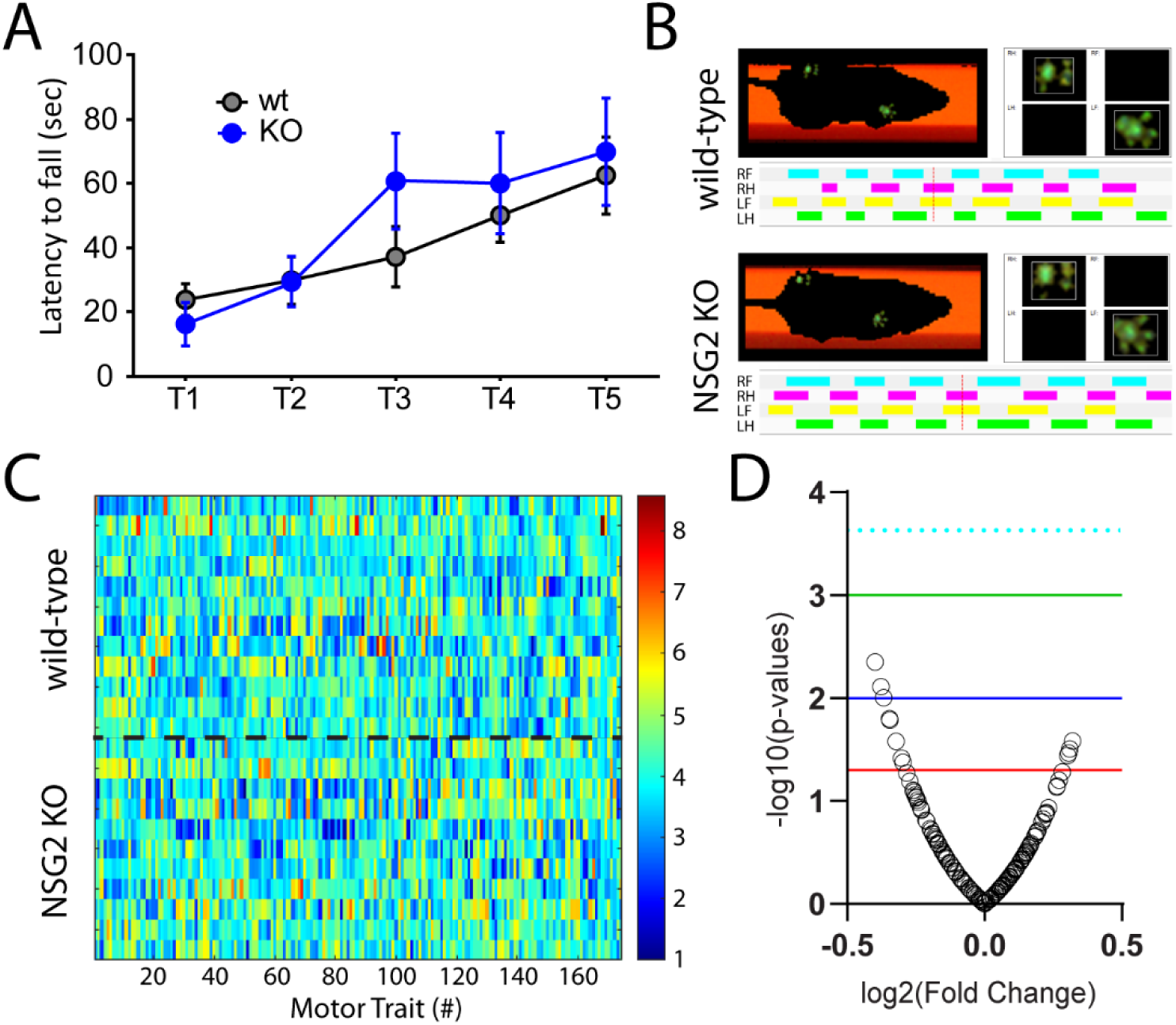
NSG2 KO animals display normal motor coordination. (A) The latency to fall from the rotating rod (4-40RPM) is presented. Means from both groups increased significantly across trials but no differences between wt and KO animals were observed. Results are expressed as mean ± SEM. (B) Representative compliant runs of wt (upper) and NSG2 KO (lower) animals depicting automated detection of right front (RF), right hind (RH), left front (LF), and left hind (LH) paws. (C) A heat map of transformed data for each trait (column) and mouse (row) were evaluated statistically and are represented as a volcano plot in (D). While several individual traits were found to differ between genotypes (red line p = 0.05), none survived False-Discovery Rate correction (dashed cyan line p = 0.0002). Blue line p = 0.01; green line p = 0.001.

### NSG2 KO animals show normal affective behavior

To test for altered affective behavior in NSG2 KO animals we used two complementary assays. We first performed the open field task to measure the mouse’s propensity to avoid exposure in open areas as well as explore a novel environment. Figure 3A shows representative heat maps for wt (upper panel) and NSG2 KO (lower panel) animals, which both illustrate relatively longer times spent in the corners and border regions compared to the center. Figure 3B illustrates pooled data showing a significant main effect of arena location (duration in border vs. center; F_(1,44)_=70.4, p<0.0001) but no significant effect of genotype (F_(1,44)_=0.02, p=0.88), or interaction (F_(1,44)_=0.92, p=0.34) for time spent in these areas. In addition, no differences were found between genotypes for total distance traveled (F_(1,44)_=1.57, p=0.22), number of entries (F_(1,44)_=0.86, p=0.36), nor velocity (F_(1,44)_=3.35, p=0.07) in either center or border regions (Supplementary Figure 1A-C). We next used the elevated zero maze, which adds elements of elevation and relatively narrow, open platforms to assess anxiety-like behavior across groups (Braun et al., 2011). Similar to the open field task, our results demonstrate a significant main effect of arm location (F_(1,44)_=102.8, p<0.0001), where animals spent significantly more time in the closed arm (p<0.0001), but no significant interaction between genotype and duration spent in either arm of the apparatus (Figure 3C-D; F_(1,44)_=2.243, p=0.14). Taken together, NSG2 KO animals showed no differences in anxiety-related behavior compared to wt animals.

**Figure 3.**
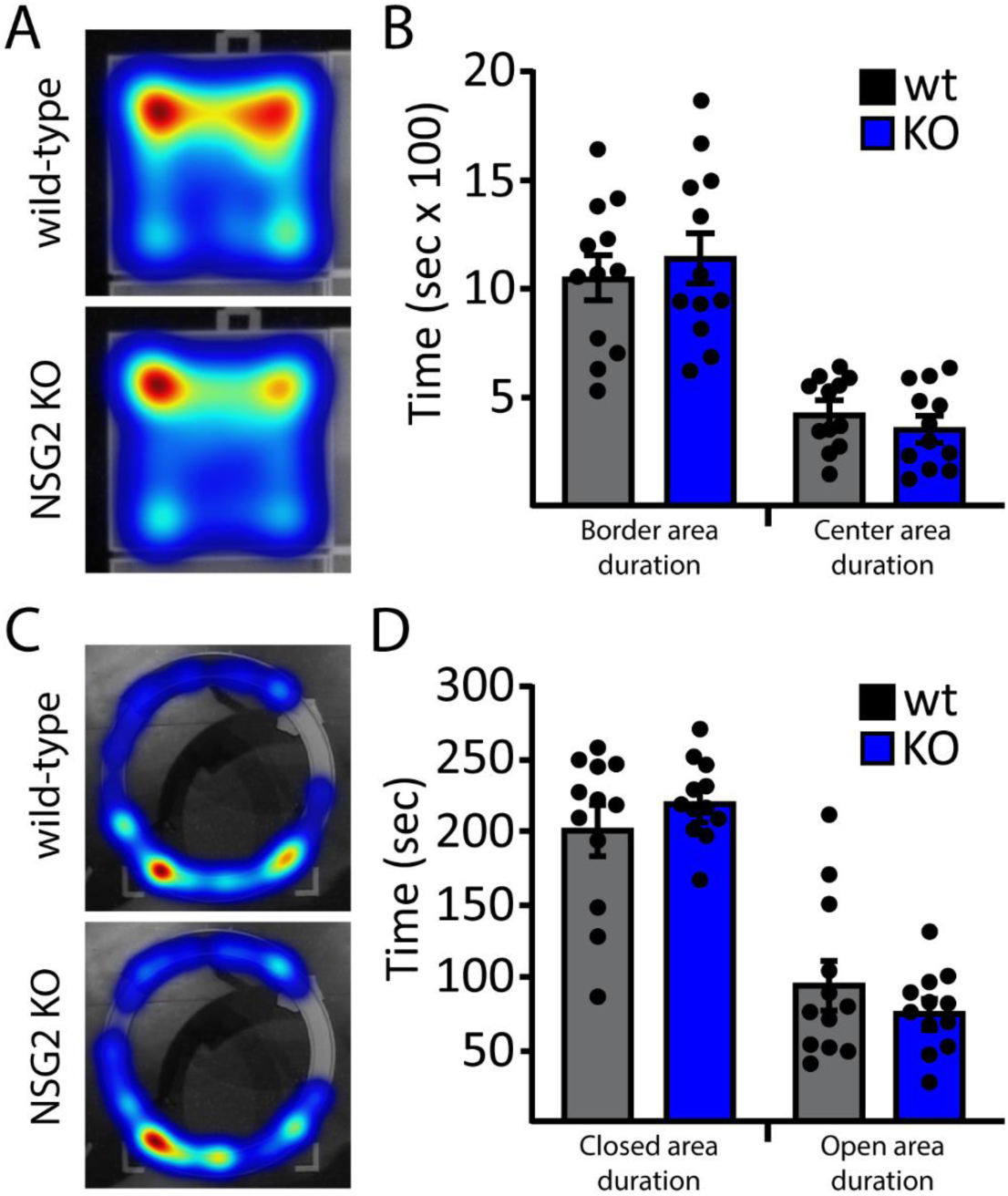
Loss of NSG2 does not cause alterations in anxiety-like behavior. (A) Representative heat maps for wt (upper panel) and NSG2 KO (lower panel) animals in open field task. (B) Pooled data reveal no significant differences in time spent along the border and in the center of the apparatus. (C) Representative heat maps for wt (upper panel) and NSG2 KO (lower panel) animals in the EZM task. (D) Pooled data reveal no significant difference between groups for time spent in the closed arms and open arms. Results are expressed as mean ± SEM. n=12/genotype.

### Loss of NSG2 decreases activity during subjective daytime

Given the relevance of disrupted sleep and circadian rhythm endophenotypes to multiple neurological and psychiatric disorders (Freeman et al., 2020; Zhang et al., 2022), we assessed the diurnal locomotor activity of NSG2 KO animals using homecage monitoring. Mice were monitored continuously over two full circadian cycles following an initial acclimation period. As predicted, a significant effect of time was observed, whereby animals displayed significantly more beam breaks during the dark cycle (active phase) compared to the light cycle (inactive phase; Figure 4A; F_(12,29)_=26.49; p<0.0001). Interestingly, we also found a significant interaction between genotype and time (Figure 4B; F_(71,1562)_=1.43; p=0.003), with loss of NSG2 causing significant reductions in activity during the wake phase compared to wt animals (p=0.004; Bonferroni’s multiple comparison test). Intriguingly, these findings are in direct opposition to those found for NSG1 KO animals, which showed significantly *greater* activity during their active phase ((Austin et al., 2022); see discussion). NSG2 KO animals showed no changes in overall activity levels during their inactive phase (Figure 4A, C; p=0.45; Bonferroni’s multiple comparison test), similar to findings for NSG1 KO animals (Austin et al., 2022).

**Figure 4.**
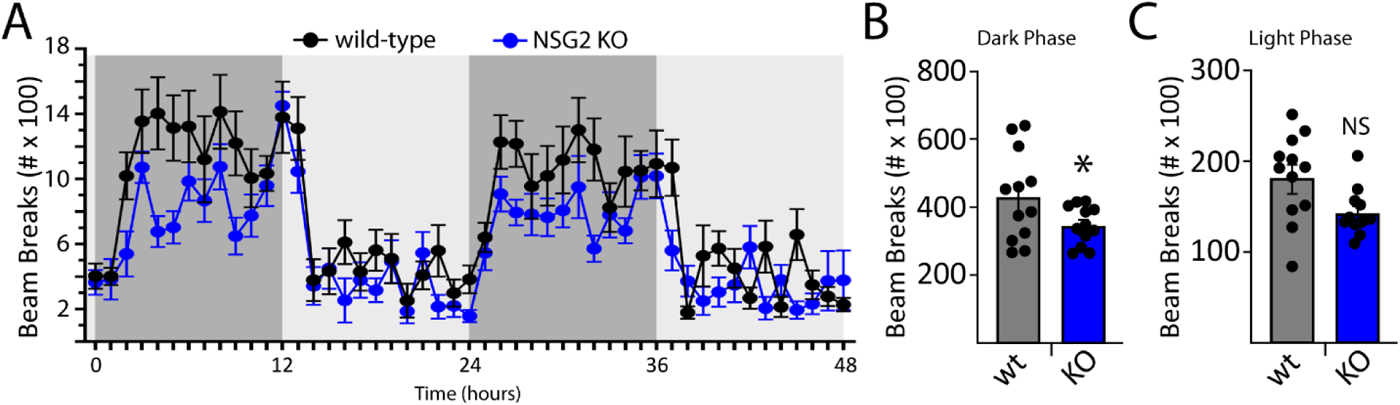
NSG2 KO selectively decreases activity during the active phase. (A) Homecage behavioral monitoring using photocell beam breaks to measure horizontal activity across 48 hours. There was a highly significant effect of phase (light vs dark) where animals showed significantly more activity during dark periods (active phase; p<0.0001). (B) NSG2 KO animals were significantly less active than wild-type animals during the active phase (p<0.05), while no differences were found for the light phase (C; inactive phase, p>0.05). Results are expressed as mean ± SEM. n=12/genotype.

### NSG2 KO enhances cue-directed learning and memory

To assess whether KO of NSG2 affects cognitive abilities we first performed a touchscreen Discrimination-Reversal (D-R) task, which relies on both frontocortical as well as dorsal striatal circuits (Izquierdo et al., 2017; Schoenbaum et al., 2007). We initially trained animals to discriminate between a target (rewarded) and non-target (punished) image (Figure 5A), where we required animals to reach an 85% correct choice criterion prior to advancing to the reversal stage. During the initial discrimination phase, NSG2 KO animals needed 27.3% fewer trials, on average, to reach criterion compared to wt animals, although this failed to reach statistical significance (Figure 5B, t_(23)_=1.461, p=0.16). NSG2 KO animals did not show a significant difference in reaction time (Supplemental Figure 3A, t_(23)_=0.795, p=0.43) but did show modest slowing of magazine latency, or time to retrieve reward during this phase (Supplemental Figure 3B, t_(23)_ = 2.668, p=0.01). Interestingly, when the target and non-target images were reversed we found a significant effect of genotype (t_(23)_ = 2.085, p = 0.048) on total trials across the entire course of the reversal problem (Figure 5C). Because early-to-mid reversal (<50% correct) is governed by orbitofrontal signaling (Marquardt et al., 2017) and late reversal (>50%) involves a shift to dorsal striatal signaling (Brigman et al., 2013; Chandrasekaran et al., 2024; Marquardt et al., 2019), we segregated the reversal into these two stages as described previously (Brigman et al., 2008; Marquardt et al., 2019). When split, we observed a significant main effect of genotype (Figure 5D (F_(1,23)_=4.676, p=0.04)). During the first phase, which constitutes primarily perseveration to the previously rewarded image, NSG2 KO animals did not demonstrate significant differences compared to wt animals (Figure 5D, “<50%”; p=0.84, Bonferroni’s multiple comparisons test). However, NSG2 KO animals needed significantly fewer trials to reach criterion during the over 50 percent phase compared to wt animals (Figure 5D, “>50”; p=0.008, Bonferroni’s multiple comparisons test). Segregating by trial type (perseverative, regressive, lose-shit, or win-stay) in the trials where mice performed above 50 percent correct, we observed a main effect of genotype (F_(1,92)_ = 22.46, p <0.0001), which was largely driven by fewer win-stay trials– the dominant trial type during late reversal (Marquardt et al., 2017); Supplemental Figure 3E; p<0.0007, Bonferroni’s multiple comparisons test). NSG2 KO animals also displayed faster reaction times during the reversal phase (Supplemental figure 3C, t_(23)_=2.192, p=0.04), but significantly slower retrieval latencies (Supplemental Figure 3D, t_(23)_=2.927, p=0.008), similar to the discrimination phase. Together, these results indicate NSG2 KO mice adapt their responding to a more efficient approach in late reversal, minimizing the number of trials to reach criterion.

**Figure 5.**
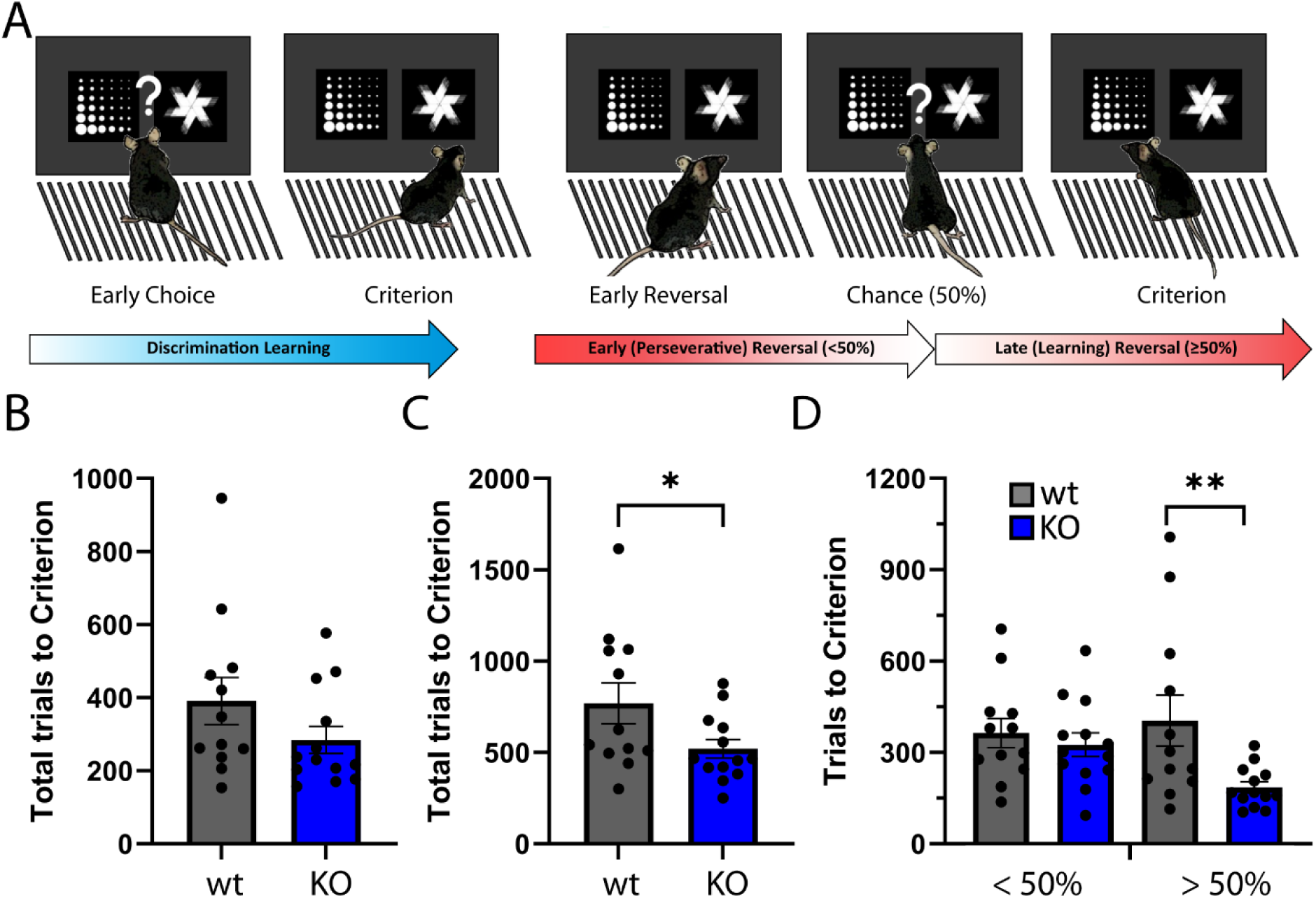
Loss of NSG2 causes enhance behavioral flexibility. (A) Schematic of touchscreen-based Discrimination-Reversal Paradigm. Animals are trained to respond to 85% correct for two consecutive sessions in the initial Discrimination before the shape choice is reversed. Early-to-chance reversal is represented as fewer than 50% correct responses, while above 50% correct until criterion (85% over two days) is considered late reversal. B. Average total trials to reach criterion in Discrimination. C. Average total trials to reach Criterion in Reversal. D. Average total trials during early-to-chance reversal (Under 50) and average total trials during late reversal (over 50). Each dot represents an individual animal. * indicates p<0.05, by two-tailed independent t-test in C and mixed-model ANOVA with Bonferroni’s multiple comparisons test in D.

To examine whether other brain regions influence altered cognitive ability, we tested animals on the Morris Water Maze (MWM; Figure 6A) which explores hippocampal-dependent spatial learning and memory (Vorhees and Williams, 2006; Whishaw et al., 1997). Both wt and NSG2 KO animals showed similar rates of acquisition during hidden platform training (Figure 6A; main effect of genotype: F_(1,22)_ = 0.001, p = 0.97), whereby both groups significantly improved from session 1 to 6 (main effect of trial (F_(3.639,80.06)_ = 13.74, p < 0.0001)) reaching criterion escape latencies under twenty seconds by day 6 on average (Figure 6A). Interestingly, despite appropriately learning the platform location, wt animals did not significantly differ from chance levels for time spent in the Target quadrant on the probe trial (Figure 6B; t_(11)_=1.229, p=0.24). However, NSG2 KO animals displayed superior retention of the task, spending significantly greater time in the Target quadrant than expected by chance (Figure 6B; t_(11)_=2.655, p=0.022). There were no differences between groups in latency to reach a visible platform (Figure 6C, Mann-Whitney *U* = 54, p=0.32) indicating no difference in motivation to escape the aversive condition.

**Figure 6.**
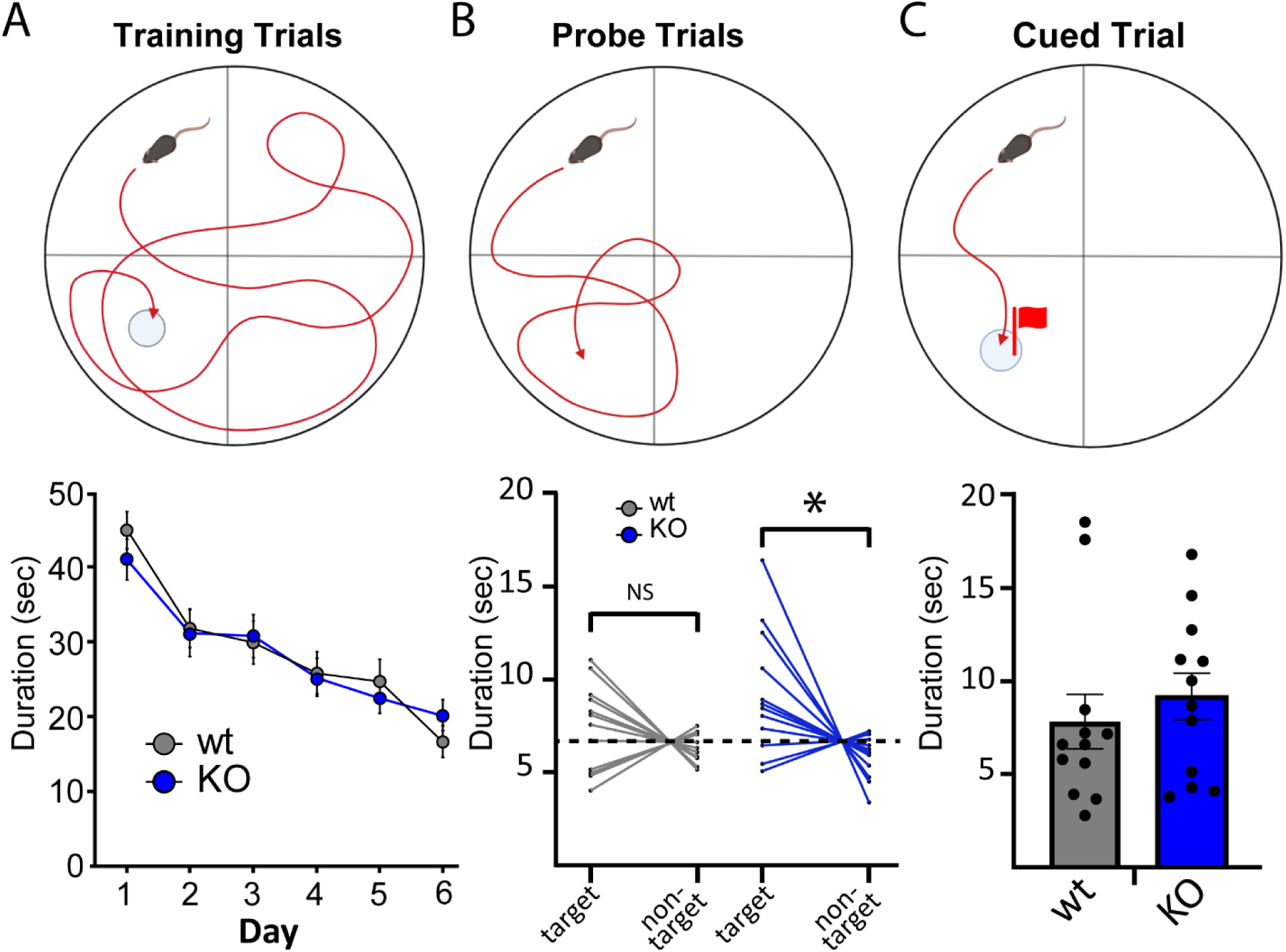
NSG KO animals display enhanced recall in Morris Water Maze. (A) Schematic of Morris Water Maze design for training (left), probe (middle), and cued (right) trials. B. Mean and SEM for latency to reach the hidden platform on each training day. C. Time spent in the target vs. non-target quadrant for each animal. Dotted line represents 25% of the total time, which is equivalent to chance. D. Mean and SEM for latency to the platform during the cued trial when the platform is made visible by a cue placed on top.

Finally, we used Trace Fear Conditioning (TFC) as a means of examining both hippocampal and amygdala-dependent associative learning and memory. Here, animals were trained to associate a tone (CS) with foot shock (US) across 5 pairings, each of which was separated by a 20 second trace period. We previously found that NSG1 KO animals showed no significant differences in the TFC task, despite significant increases in anxiety-like behavior (Austin et al., 2022). In stark contrast, we found that loss of NSG2 resulted in a significant acceleration of acquisition of CS-US pairing, as illustrated by increased freezing behavior during both the tone (main effect of genotype: F_(1,22)_ = 16.43, p = 0.0005) and trace periods (main effect of genotype: F_(1,22)_ = 8.85, p = 0.007; Figure 7A-B, respectively). Furthermore, NSG2 KO animals demonstrated significantly increased freezing behavior during the tone 24 hours after training, when no US (shock) was present (Figure 7C,t_(22)_ = 2.69 p=0.01). No significant difference between groups was found during the trace period on day 2 of testing, but this was likely due to the high level of freezing in the wt animals (Figure 7D, t_(22)_ = 1.277p=0.21). Taken together, results from cognitive tests suggest that loss of NSG2 causes accelerated learning and enhances retention across multiple brain regions including dorsal striatum, hippocampus and amygdala.

**Figure 7.**
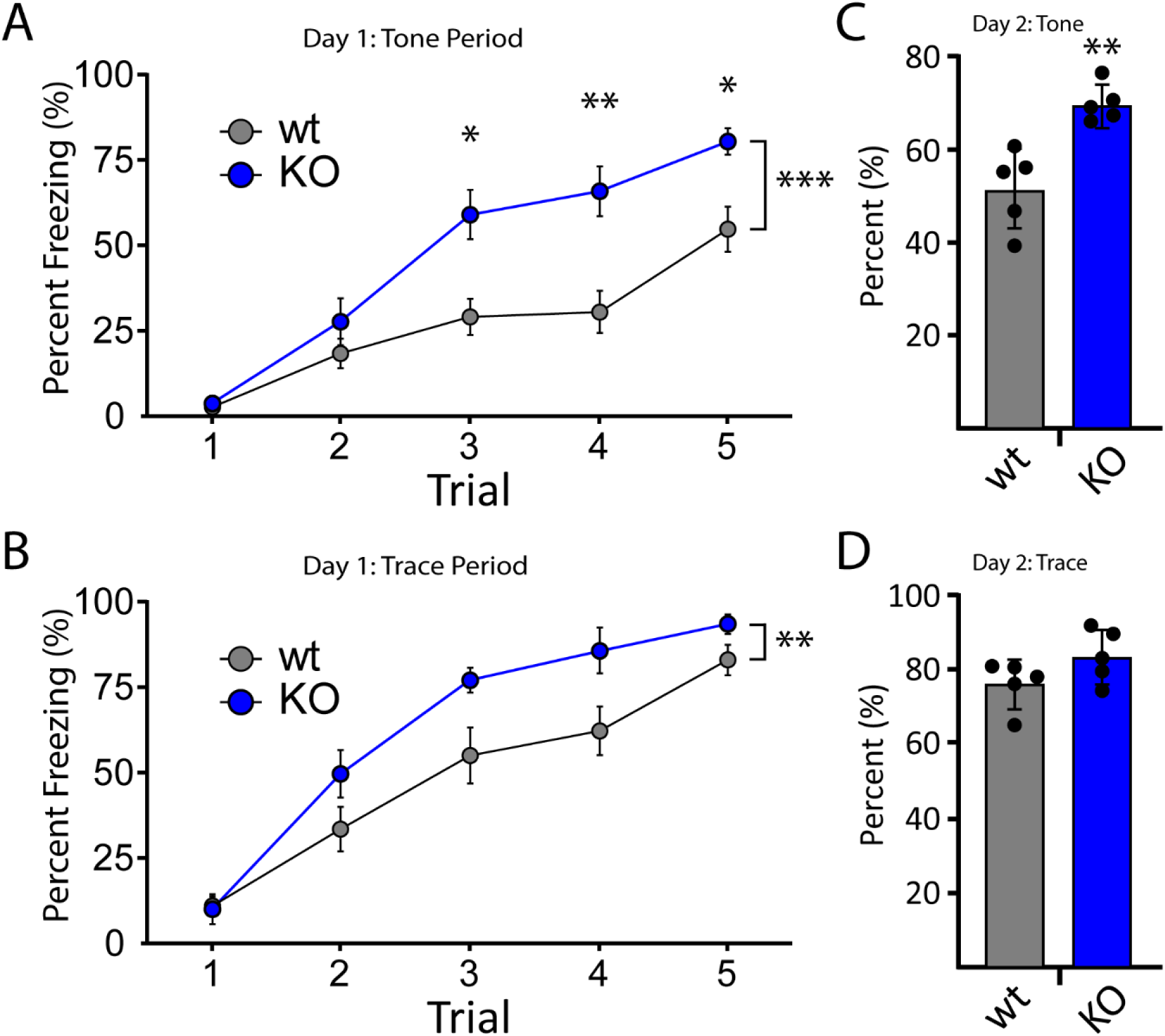
NSG2 KO animals display accelerated fear learning and augmented associative memory. (A-B) Percentage of time spent freezing during the tone period (A) and trace period (B) is illustrated across five acquisition trials for both wt and NSG2 KO mice. Compared to wt mice, NSG2 mutants exhibited significantly increased freezing times during the training period (p<0.02). (C) NSG2 KO animals also displayed greater freezing behavior during the tone period (p<0.05) when presented in the same chamber 24 hours following training but without footshock stimuli. (D) No differences were observed between groups during the trace period when presented in the same chamber 24 hours following training but without footshock stimuli (p>0.05). Results are expressed as mean ± SEM. n=12/genotype.

## DISCUSSION

A proposed model for NSG protein function during synaptic plasticity suggests a primary role of NSG1 in LTP, via sorting internalized AMPARs for recycling back to the plasma membrane following induction (Muthusamy et al., 2015). In contrast, the proposed function of NSG3 (Calcyon) lies in regulating LTD by promoting internalization of surface AMPARs via clathrin-mediated endocytosis (Davidson et al., 2009; Xiao et al., 2006). Currently, there are no data regarding the role of NSG2 during plasticity-inducing paradigms. However, under basal transmission our data suggest that NSG2 promotes AMPAR surface expression, where overexpression led to increased amplitude of miniature excitatory postsynaptic currents (mEPSCs), while CRISPR-mediated KO of NSG2 reduced mEPSC frequency (Chander et al., 2019). Thus, it could be hypothesized that the role of NSG2 may be somewhat redundant to that of NSG1, whereby both proteins promote synaptic strengthening via delivery of AMPARs to post-synaptic densities (Alberi et al., 2005; Steiner et al., 2005). In contrast, the current model predicts that both NSG1 and NSG2 function in a largely complementary fashion to NSG3.

However, animal studies suggest this reductionist view is incomplete based on behavioral analyses. While knockout of NSG1 did impact motor coordination and anxiety, NSG1 KO animals showed no changes in learning and memory (Austin et al., 2022), despite its published role in synaptic plasticity. Further, NSG3 overexpression (OE) did not affect acquisition of basic associative learning nor short-term reference memory (Austin et al., 2022; Vazdarjanova et al., 2011). In contrast, loss of NSG2 caused significant acceleration of association learning, increased recall, as well and enhanced behavioral flexibility (Figures 5-7, this study). These data suggest NSG2 has a unique role in synaptic plasticity of memory circuits. Interestingly, NSG3 OE mice did show significant perseveration in both MWM reversal phase and Fear Extinction paradigms, consistent with impaired long-term depression (Davidson et al., 2009; Vazdarjanova et al., 2011). Thus, despite the difficulties is extrapolating data from overexpression studies, it is possible that in cognitive domains, NSG2 and NSG3 may have opposing effects on synaptic plasticity at least in hippocampus. As envisaged, endogenous levels of NSG2 and NSG3 provide a balance of exo- and endocytosis of AMPARs in post-synaptic membranes, respectively (Chander et al., 2019; Davidson et al., 2009; Xiao et al., 2006). This balance may provide a limit to both the number of AMPARs that can be added and removed from synapses, resulting in limits to both potentiation or depression. Obviously, the major limitations to the comparison across these studies is the lack of behavioral analysis of NSG3 KO animals as well as differences across some of the behavioral assays performed. In addition, future studies will need to determine whether NSG2 KO animals display enhanced LTP as well as perseveration on hippocampal-dependent tasks, as enhanced fear responding may be maladaptive. However, taken together, while NSG1 appears critical in motor and anxiety-related pathways, NSG2 and NSG3 may be primarily involved in associative learning and memory via differential AMPAR trafficking.

Multiple transgenic (Tg) and KO mouse models demonstrate enhanced learning abilities, but the previously associated mechanisms underlying behavioral changes converge on a small number of known plasticity pathways, and AMPAR trafficking has yet to be implicated (Lee, 2015). For example, increasing NMDA receptor activity either directly by overexpression of the NR2B subunit itself (Cui et al., 2011; Tang et al., 1999), indirectly by promoting delivery of NR2B to synapses by KIF17 (Wong et al., 2002), or via changes to various signaling factors such as p25 (Fischer et al., 2005), Cdk5 (Hawasli et al., 2007) or ORL1 (Mamiya et al., 2003; Manabe et al., 1998), cause augmented performance in a variety of hippocampal- and amygdala-dependent tasks as well as enhanced synaptic plasticity. Similar results have been found when targeting downstream signaling pathways that converge on translation- and CREB-dependent gene expression (Chen et al., 2003; Hoeffer et al., 2008; Khoutorsky et al., 2013; Suzuki et al., 2011; Tsai et al., 2012) as well as gene changes that have a net effect of decreasing inhibition via GABA receptors and potassium channels (Miyake et al., 2009; Moore et al., 2010; Murphy et al., 2004; Zhu et al., 2011). AMPARs play an essential role in synaptic plasticity and are a pharmacological target for cognitive enhancement in clinical trials (Partin, 2015). However, Tg and KO animals where genes primarily involved in AMPAR trafficking and function are targeted, result in either no change, or significant *deficits* to learning and memory. For instance, single deletions of most Transmembrane AMPAR regulatory proteins (TARPs) do not show obvious behavioral phenotypes (Letts et al., 2005; Menuz et al., 2008; Milstein and Nicoll, 2009; Rouach et al., 2005), whereas mutations of TARP2 cause learning and memory deficits (Caldeira et al., 2022). Deficits of learning and memory are also observed in knockout animals of AMPAR binding proteins GRIP1 (Tan et al., 2020), GRASP1 (Chiu et al., 2017), LARGE1 (Seo et al., 2018), as well as the PICK1-associated protein ICA69 (Chiu et al., 2023). Even in cases where modifications to AMPARs cause enhanced LTP, learning and memory deficits are observed (Guntupalli et al., 2023).

Previous studies implicate NSG2 in binding, trafficking, and surface expression of AMPARs (Chander et al., 2019), but NSGs play a role in a plethora of additional cellular functions. Thus, deciphering the mechanism by which loss of NSG2 enhances learning and memory may provide unique insights into novel mechanisms for the regulation of synaptic plasticity. While a large body of literature implicates recycling endosomes in supporting synaptic function via AMPAR trafficking (Esteves da Silva et al., 2015), NSG2 is primarily localized to late endosomes (LEs), with a small proportion contained within early endosomes and at the plasma membrane (Yap et al., 2017). A small number of intriguing reports have suggested that acidic vesicles within dendrites support structural plasticity of dendritic spines via exocytosis of matrix metalloproteases to degrade extracellular matrix (Goo et al., 2017; Grochowska et al., 2023; Padamsey et al., 2017). Furthermore, multi-vesicular bodies (MVBs) containing undergo fusion with the plasma membrane to release extracellular vesicles (Hessvik and Llorente, 2018), and NSG1 has been localized to intraluminal vesicles (Utvik et al., 2009). Thus, it is possible that NSG2 plays a role in the regulation of exocytosis of proteins/vesicles to peri-synaptic sites via MVBs or acidic vesicles that are involved with relatively novel forms of plasticity. Alternatively, NSG2 may regulate trafficking or post-translational processing of other types of receptors and/or signaling pathways. NSG3 regulates the dopamine D1 receptor cycling between plasma membrane and endosomal compartments (Ha et al., 2012). Finally, NSG1 and NSG3 regulate proteolytic processing of APP and Neuregulin-1, respectively, proteins known to alter synaptic function. Future studies are required to determine whether loss of NSG2 converges on standard mechanisms of synaptic function, or whether novel mechanisms may underlie enhanced cognition in NSG2 KO animals.

## DATA AVAILABILITY STATEMENT

The data that support the findings of this study are available from the corresponding author upon reasonable request.

## ACKNOWLEDGMENTS

We would like to thank members of the UNM-HSC Preclinical Core (Drs. LeeAnna Cunningham and Carissa Milliken) of the Center for Brain Repair and Recovery for their training and assistance with all the behavioral experiments and for assistance with brain section imaging.

## MATERIALS AND METHODS

All experimental procedures adhered to the US Public Health Service policy on humane care and use of laboratory animals, and were approved by the Institutional Animal Care and Use Committee at the University of New Mexico Health Sciences Center. In addition, experimenters were blinded to group assignments of all animals for the experiments described.

### *NSG2* mutant mouse model

Constitutive “NSG2 KO em1” (C57BL/6N-Nsg2^em1(IMPC)KMPC/KMPC^) mice were purchased from and generated by the Korean Mouse Phenotyping Center (KMPC), and details of the CRISPR-mediated gene targeting strategy can be found at https://mousephenotype.kr/. Heterozygote (NSG2^+/-^) animals were bred to produce a viable colony from which the cohort reported here was generated. Offspring were weaned at between 21-23 days of age, ear tagged, and housed with same sex littermates at 22°C on a 12-h reverse light/dark cycle (dark period: 0800-2000). The mice had free access to standard chow and water at all times except where noted below (D-R learning paradigm). Behavior tests were performed on a cohort of 12 wild-type (wt; NSG2^+/+^), and 13 KO (NSG2^-/-^) male mice (total n=25) and all animals were between the ages of 2 and 6 months during testing. NSG2 genotyping was performed via Polymerase Chain Reaction (PCR). Tail snips (∼0.5 cm) were digested in 200 μl of DirectPCR Lysis Reagent (Viagen Biotech, Los Angeles, CA) and 2μl of Proteinase K (Zymo Research, Irvine, CA) at 55°C for 24h followed by incubation at 95°C for 10min to inactivate Proteinase K. PCR amplification of NSG2 was carried out using Q5 DNA polymerase according to the manufacturer’s recommendations (New England Biolabs, Ipswich, MA). The following primers were used to determine specific allelic expression: 5’-AGGGGGCTTGTCATCTCTGA-3’ (*NSG2* forward), 5’-GTCCCCTTCCATTCCATCCC-3’ (*NSG2* reverse). 1 μl of digested genomic DNA was used in a 20 μl PCR reaction containing both primers (5 μM each). The following cycling parameters were used: 1 cycle of 95°C for 3 min, 30 cycles of 95°C (30s), 68°C (30s), 72°C (30s) and 1 cycle of 72°C for 5 min (C1000 Touch Thermal Cycler, Bio-Rad, Hercules, CA). 2–5 μl of the PCR reactions were subjected to gel electrophoresis using a 3% agarose gel in tris-borate-ethylenediamine tetraacetic acid buffer. Amplicons were visualized to identify mice with wt (227 bp), KO (220 bp), and heterozygote (227 and 220 bp) genotypes.

### Western Blot Analysis

Brains from wt and NSG2 KO mice were homogenized in 1X RIPA buffer (150 mM sodium chloride, 1.0% Triton X-100, 0.5% sodium deoxycholate, 0.1% SDS, 50 mM Tris, pH 8.0), 1x proteinase inhibitor (Thermo, #A32955) with a micro-homogenizer while kept on ice. Detergent-insoluble material was removed by centrifugation at 13,000x g for 10 min. Protein concentration was determined using the Pierce BCA Protein Assay Kit (#23225). Twenty micrograms of protein were mixed with 1X NuPAGE LDS Sample Buffer (Invitrogen, #NP0007), 1X NuPAGE Sample Reducing Agent (Invitrogen, #NP0009) and incubated at 95°C for 20 minutes. The samples were separated on a 15% Tris-Glycine gel with a 4% stacker using a Tris/Glycine/SDS buffer and transferred overnight onto an Immobilon-FL Transfer Membrane using a Mini Trans-Blot Cell (Bio-Rad). Membranes were blocked for one hour (0.5% Non-Fat Dry Milk in PBS) and then probed with the primary antibody: rabbit anti-NSG2 (Abcam; ab189513, 1:1000) for 2h at room temp (0.5% Non-Fat Dry Milk/0.1% Tween-20 in PBS). The blots were washed 3x5min with PBS and incubated one hour with secondary antibody [Goat anti-Rabbit horseradish peroxidase (HRP) conjugate (Jackson ImmunoResearch Laboratories, Inc., 111-035-003, 1:10,000)]. After another 3x5min wash with PBS, signal was developed using SuperSignal West Pico PLUS Chemiluminescent Substrate (Thermo, #34580), and imaged on a Bio-Rad ChemiDoc. The unstripped membrane was then probed with the primary antibody: mouse anti-NSG1 (Santa Cruz; sc-390654, 1:1000) and secondary: Goat anti-Mouse horseradish peroxidase (HRP) conjugate (Jackson ImmunoResearch Laboratories, Inc., 115-035-003, 1:10,000).

### Immunohistochemistry

WT and NSG2 KO mice were anesthetized and perfused with phosphate buffered saline (PBS) followed by 4% paraformaldehyde (PFA). Brains were removed and immersed in 4% PFA, 20% sucrose, and 30% sucrose, each for 12–24 h. Sagittal sections were sliced on a sliding knife microtome (American Optical). Free-floating sections were permeabilized and blocked simultaneously in 0.5% Triton X-100 (Sigma-Aldrich, St. Louis, MO) and 10% donkey serum (MilliporeSigma) for 1 hr in PBS. Sections were stained with goat anti-NSG1 (Thermofisher; PA5-37939, 1:1000) and rabbit anti-NSG2 (Abcam; ab189513, 1:500) in 0.25% Triton and 5% donkey serum in PBS overnight at 4°C. Sections were washed three times in PBS followed by secondary antibody labeling (donkey anti-goat and donkey anti-rabbit [1:1000]; Thermofisher) in the same buffer as primary antibodies. Sections were then mounted to microscopy slides (Superfrost plus, Fisher Scientific), immersed in Fluoromount-G, and imaged on a slide-scanning microscope (Zeiss Axion Scan.Z1) with Colibri 7 LED light source. Standard fluorescence calibration was performed on wild-type control brain sections to ensure proper dynamic range of signals and imaging conditions were maintained on NSG2 KO brains

### Behavioral Analyses

All behavioral assays were performed essentially as described previously (Austin et al., 2022) with the exception of the Morris Water Maze and touchscreen discrimination-reversal tasks. Where appropriate, mice were acclimated to testing rooms for at least 30 min. prior to testing and all apparati were cleaned using 70% ethanol and dried between individual trials to eliminate olfactory distractions. All of the experiments were performed during the active phase between the hours of 0900–1200, except the Morris Water Maze, which was separated into two session (0930-1100 and 1200-1430). Tests were performed in a room illuminated by red lights, except for homecage behavioral monitoring, which occurred during both light and dark phases. Behavioral assays were performed in the following order: discrimination-reversal operant learning, open field, rotarod, Catwalk XT, elevated zero maze (EZM), trace fear conditioning (TFC), homecage monitoring. All tests were separated by a 24–48 interval.

### Anxiety-related tasks

The <u>open-field test</u> was used to assess exploratory behavior, anxiety, and gross locomotion. Mice were randomly placed in one of the corners of a white Plexiglas open field chamber (29 X 29 X 29cm) and allowed to freely explore for 30 minutes. An overhead camera (Med Associates Basler acA1300-60) and Ethovision tracking software XT (Noldus Information Technology, Wageningen, Netherlands) were used to track velocity, distance traveled, and duration in the center or the border areas. The <u>elevated zero maze</u> <u>(EZM)</u> consisted of an elevated circular platform (64 cm high; 50.8 cm min diameter and 60.96 cm max diameter) with two opposing enclosed sections enclosed by a wall (20 cm high) and two opposing open sections, each with a platform 5 cm wide. Each mouse was randomly placed in one quadrant within the maze and allowed to explore for 5 min while tracked via the same overhead camera and video-tracking software as noted above to monitor the position of the central point of the mouse body. The software measured latency and cumulative time spent in the open and closed areas of the maze.

### Motor-related tasks

The <u>rotarod task</u> was used to assess motor performance and fatigue resistance in rodents using the Panlab Rota Rod model LE8205 (Harvard Apparatus, Holliston, MA). The rod was initiated to rotate at a constant initial speed of 4 RPM. Once the mouse was positioned, the rod accelerated to 40 RPM in 300 s. The time spent on the rod and the RPM reached at the time of falling was captured for each of five trials. A 30-s inter-trial interval (ITI) was used throughout the experiment. The <u>CatwalkXT</u> <u>system</u> (Noldus, Leesburg, VA) was used to analyze gait and fine motor coordination. Mice were allowed to walk freely across the glass walkway to reach a dark goal box and footprint detection occurred as animals passed through a black tunnel illuminated from one side by a reflected green fluorescent light. Three trials capturing at least three stride lengths each were captured for each animal during one testing session. Trials were included in the analysis (were “compliant”) if the mouse crossed the recording area in under 5 seconds and did not show a maximum speed variation greater than 60%. Compliant trials were analyzed via automatic detection but were also reviewed manually; trials that did not represent continuous forward movement were excluded. The extraction of 174 traits was performed in an automated fashion using Catwalk XT 8.1 Software. These data were then exported and evaluated statistically as described previously (Jacquez et al., 2021). Briefly, data for each trait was z-score transformed and normalized, and then evaluated statistically for group differences using FDR-corrected unpaired T-tests.

### Circadian regulation

Spontaneous <u>homecage activity</u> was collected in wild-type and NSG2 KO mice to assess diurnal cycles and locomotor behavior in a non-aversive environment. All mice were individually housed in a standard home cage with corncob bedding with ad libitum food and water and left undisturbed for a 72-h period under their normal illumination conditions (lights on at 2000h and off at 0800h). Horizontal activity was automatically measured by photocell beam break for 72 h using the PAS-Homecage system (San Diego Instruments, San Diego, CA). The first 9h were considered an acclimation period and excluded from analysis, while the middle 48h (two cycles) were used for data analysis (starting at 0800h, light offset). Repeated measures ANOVA was performed across the entire circadian period and two-way ANOVA was performed on cumulative light and dark activity across the 48h period to assess light (sleep) and dark (wake) phases separately.

### Learning and memory

Pavlovian <u>trace fear conditioning</u> (<u>TFC</u>) was used to measure associative learning. Animals were first placed into a Habitest® System (Coulbourn Instruments, Allentown, PA) for 180 s to habituate to apparatus. After the no stimulus period, a 90 dB, 5000 Hz white noise auditory tone was presented for 30 s (conditioned stimulus [CS]) followed by a 20-s interval that terminated with a 2 second 0.7 mA footshock (unconditioned stimulus [US]). After the first CS/US pairing there was a 90-s interval followed by another CS/US pairing. The CS/US pairings were repeated 5 times on day 1 (acquisition) with a randomized interval averaging 120 s. Each session was ended 60 s after the final pairing. Memory for the CS was tested absent the shock in the same chamber 24 h (day 2) after acquisition. <u>Morris Water Maze</u> (MWM): Equal numbers of wt and KO animals were separated into two sessions (mid-morning/mid-afternoon) to minimize inter-trial interval between animals. All animals were trained for 6 days (4x1min trials/day, 15 min inter-trial interval) to find a hidden platform submerged beneath opaque (white, non-toxic paint) water (26°C) using spatial cues on the wall, as previously described (Vorhees and Williams, 2006). Platform location by quadrant as well as animal placement were randomized to eliminate bias. Animals were singly housed and kept warm in cages between trials. On the seventh day, a probe trial was performed via removal of the platform and the percentage of time spent in the target quadrant vs. the other quadrants was recorded as a measurement of spatial working memory. On day 8, animals were tested on a single cued trial where a tall, visible cue was placed on the platform in the target quadrant. Time to reach the platform was recorded as a measure of general motor function. All behavioral data were collected using Ethovision XT8 software (Noldus, Netherlands), and then processed in Prism (GraphPad, V9).

<u>Touch screen discrimination reversal</u> (<u>D-R</u>): Operant behavior was conducted as previously described (Zimmerman et al., 2020) in a chamber measuring 21.6 x 17.8 x 12.7 cm (model # ENV-307W, Med Associates, St. Albans, VT) housed within a sound- and light-attenuating box (Med Associates, St. Albans, VT). A solid acrylic plate was used to cover the grid floor of the chamber to facilitate ambulation. A peristaltic pump delivered 10µl of liquid strawberry milkshake (strawberry Nesquik mixed with skim milk) into a magazine. A house-light, tone generator and an ultra-sensitive lever was located on one end of the chamber, while a touch-sensitive screen (Conclusive Solutions, Sawbridgeworth, U.K.) was on the opposite side of the chamber covered by a black acrylic aperture plate, which creates two 7.5 x 7.5 cm touch areas separated by 1cm and located at a height of 0.8cm from the floor of the chamber. KLimbic Software Package v1.20.2 (Conclusive Solutions) controlled and recorded stimulus presentation and touches in the response windows. Eight week-old mice were food restricted and maintained at 85% free-feeding body weight and subjected to 3 days of acclimation to the behavior room and liquid reward. Mice were habituated to the operant chamber and retrieving reward from the magazine by being placed in the chamber for ≤30 minutes with liquid available in the magazine. Once a mouse retrieved at least 10 liquid reward retrievals, during a habituation session, it began the pre-training regimen. First, mice were trained to obtain reward by pressing a lever within the chamber on an FR1 schedule. Once a mouse showed willingness to press the lever and collect 30 rewards in a <30 minute-session, it was moved to touch training. During this stage, a lever press led to the presentation of a white (variously-shaped) stimulus in 1 of the 2 response windows (spatially pseudorandomized). The stimulus remained on the screen until a response was made. Touches in the blank response window had no effect, while a touch to the white stimulus window resulted in reward delivery, immediately cued by a tone and illumination of the magazine. Once a mouse was able to initiate, touch and retrieve 30 rewards in a <30 minute-session, it was moved to the final stage of pre-training. This stage was identical to touch-training except that responses at a blank window during stimulus presentation now produced a 10-second timeout, immediately signaled by illumination of the house light, to discourage indiscriminate screen responding. Errors made on this pre-training stage (as well as on discrimination and reversal, see below), were followed by correction trials in which the same stimulus and left/right position was presented until a correct response was made. Once a mouse was able to make ≥75% (excluding correction trials) of responses at a stimulus-containing window in a 30-trial session, it was moved onto discrimination testing. Pairwise discrimination and reversal was tested as previously described (Brigman et al., 2013). Mice were first trained to discriminate two novel, approximately equally-luminescent stimuli, presented in a spatially pseudorandomized manner, over 30-trial sessions (5-second inter-trial interval). The stimulus designated as correct was counterbalanced across mice and genotypes. Responses at the correct stimulus window resulted in a single liquid food reward, cued by a 1-second tone and illumination of the magazine. Responses at the incorrect stimulus window resulted in a forced timeout, signaled by illumination of the house-light. Correction trials following errors were presented, with the same stimuli, in the same spatial orientation, until a correct response was made. Discrimination criterion was ≥85% correct responding out of 30 trials, excluding correction trials, over two consecutive sessions. Reversal training began on the session after discrimination criterion was attained. Here, the designation of correct verses incorrect stimuli was reversed for each mouse. As for discrimination, there were 30-trial daily sessions until the mice reached a criterion of ≥85% correct responding (excluding correction trials) over two consecutive sessions. For discrimination and reversal, the dependent variables were correct and incorrect trials, correction errors, reaction time (time from lever press initiation to screen touch) and magazine latency (time from screen touch to reward retrieval). In order to examine distinct phases of reversal (early perseverative and late learning), we separately analyzed errors and correction errors for sessions where performance was below 50% correct and performance above 50% correct as previously described (Brigman et al., 2008). Further analyses of trial types were performed for trials >50% correct to assess trial-type biases. These trials were separated according to consecutive trial pairs as: perseverative (incorrect-incorrect), regressive (correct-incorrect), lose-shift (incorrect-correct), win-stay (correct-correct.)

### Statistical analysis

For Homecage, TFC and rotarod analyses, we used repeated measures ANOVA with post-hoc tests unless otherwise stated. For open field and EZM, unpaired t-tests were used with statistical significance set a priori at p = 0.05 following validation of normally distributed data. Mann-Whitney U test was used for non-parametric data.

---

**Supplemental Figure 1.**
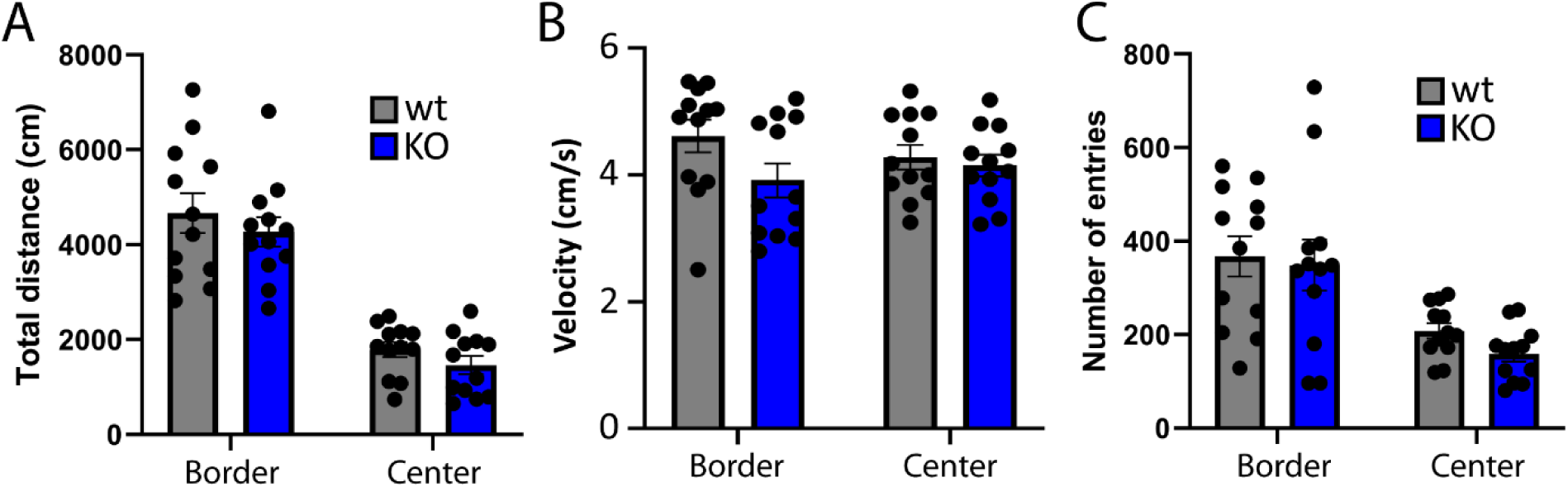
Related to Figure 3. A. Number of entries into the Border or Center areas of the Open Field box. B. Velocity in each area of the Open Field box. C. Distance traveled in each area of the Open Field box. Data are presented as mean and SEM with individual animals represented by each dot.

**Supplementary Figure 2.**
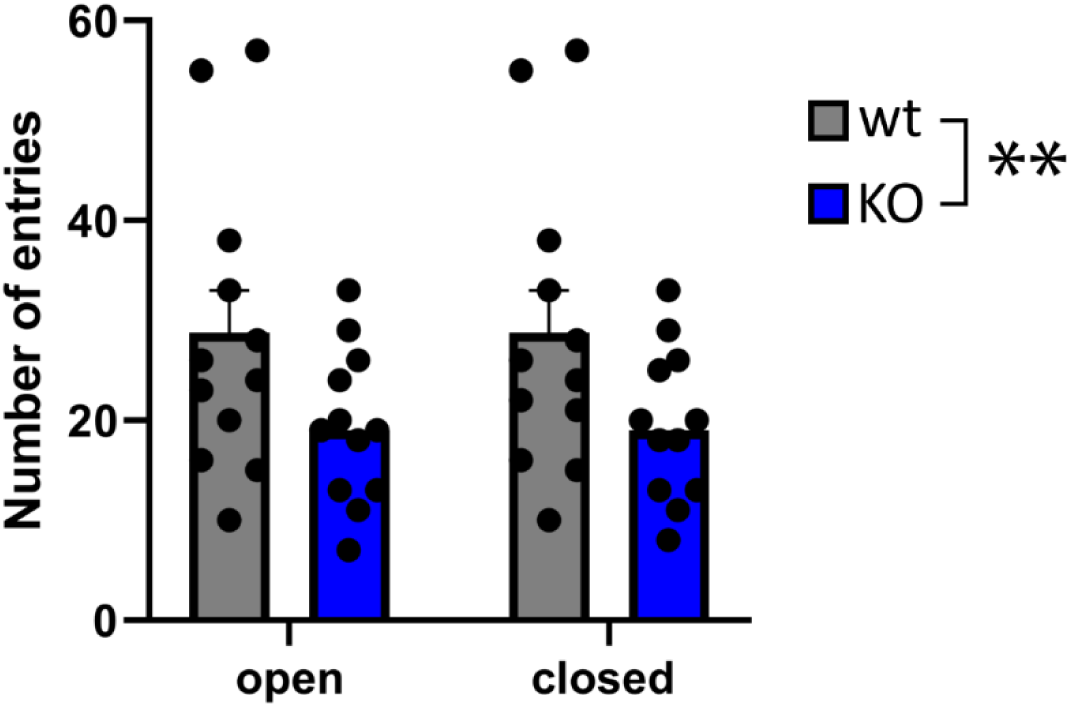
Related to Figure 3. Number of entries into each arm of the Elevated Zero Maze. Data are presented as mean and SEM with individual animals represented by each dot.

**Supplementary Figure 3.**
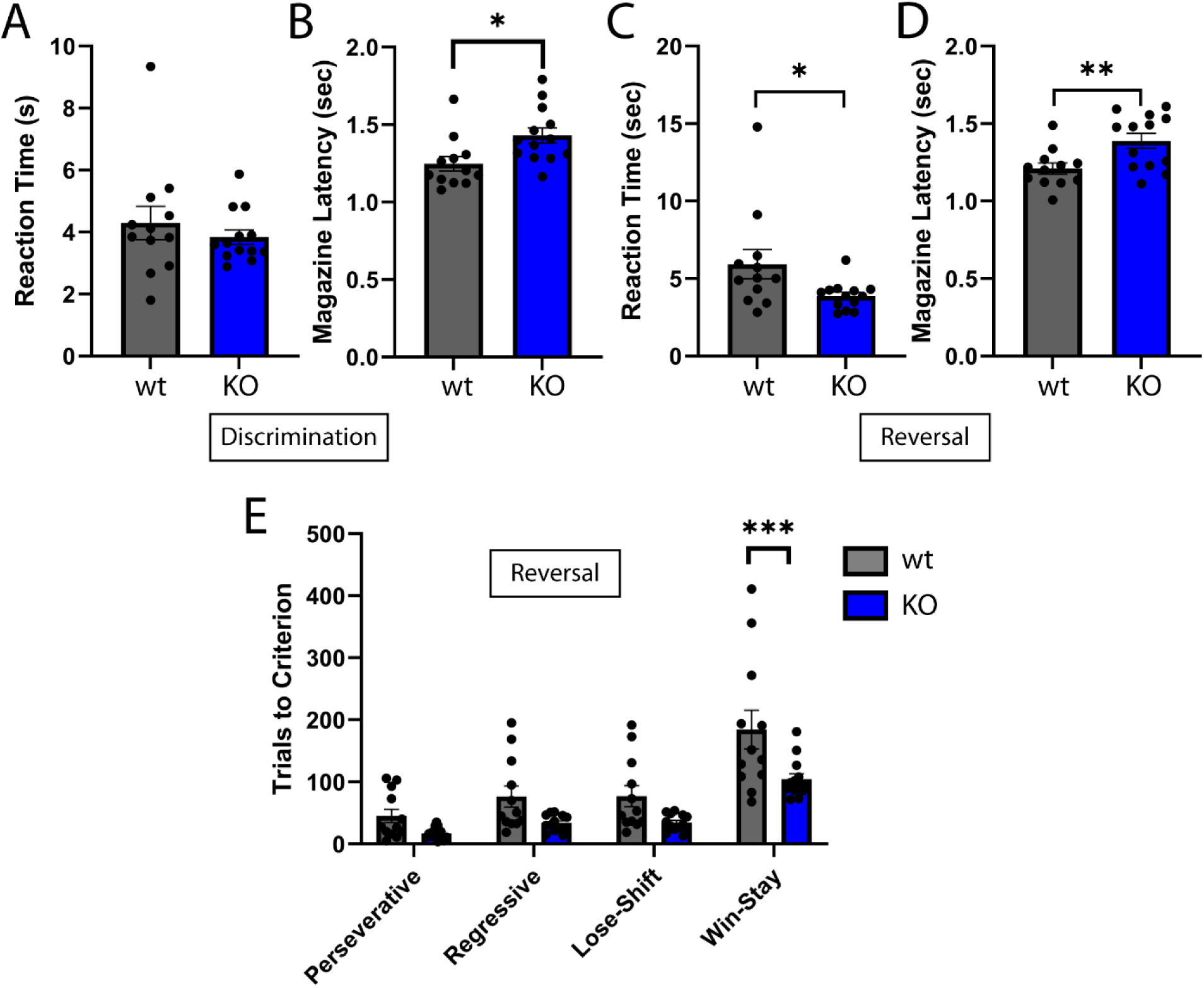
Reaction time (A) and magazine latency (B) during the initial Discrimination on the touchscreen D-R task. Reaction time (C) and magazine latency (D) during Reversal on the touchscreen D-R task. E. Average number of trials for each sub-trial category including perseverative, regressive, lose-shift and win-stay during the late (>50% correct) Reversal stage of the touchscreen D-R task. Data are presented as mean and SEM with individual animals represented by each dot. *p-value <0.05, **<0.01 by two-tailed independent t-test. In E, ***p-value <0.001 represents Bonferroni’s multiple comparisons following 2-way ANOVA.

